# Bedside formulation of a personalized multi-neoantigen vaccine against mammary carcinoma

**DOI:** 10.1101/2021.04.24.440778

**Authors:** Mona O. Mohsen, Daniel E. Speiser, Justine Michaux, HuiSong Pak, Brian J. Stevenson, Monique Vogel, Varghese P. Inchakalody, Simone de Brot, George Coukos, Said Dermime, Michal Bassani-Sternberg, Martin F. Bachmann

## Abstract

Graphical abstract
Individualized neoantigen vaccination against mammary carcinoma

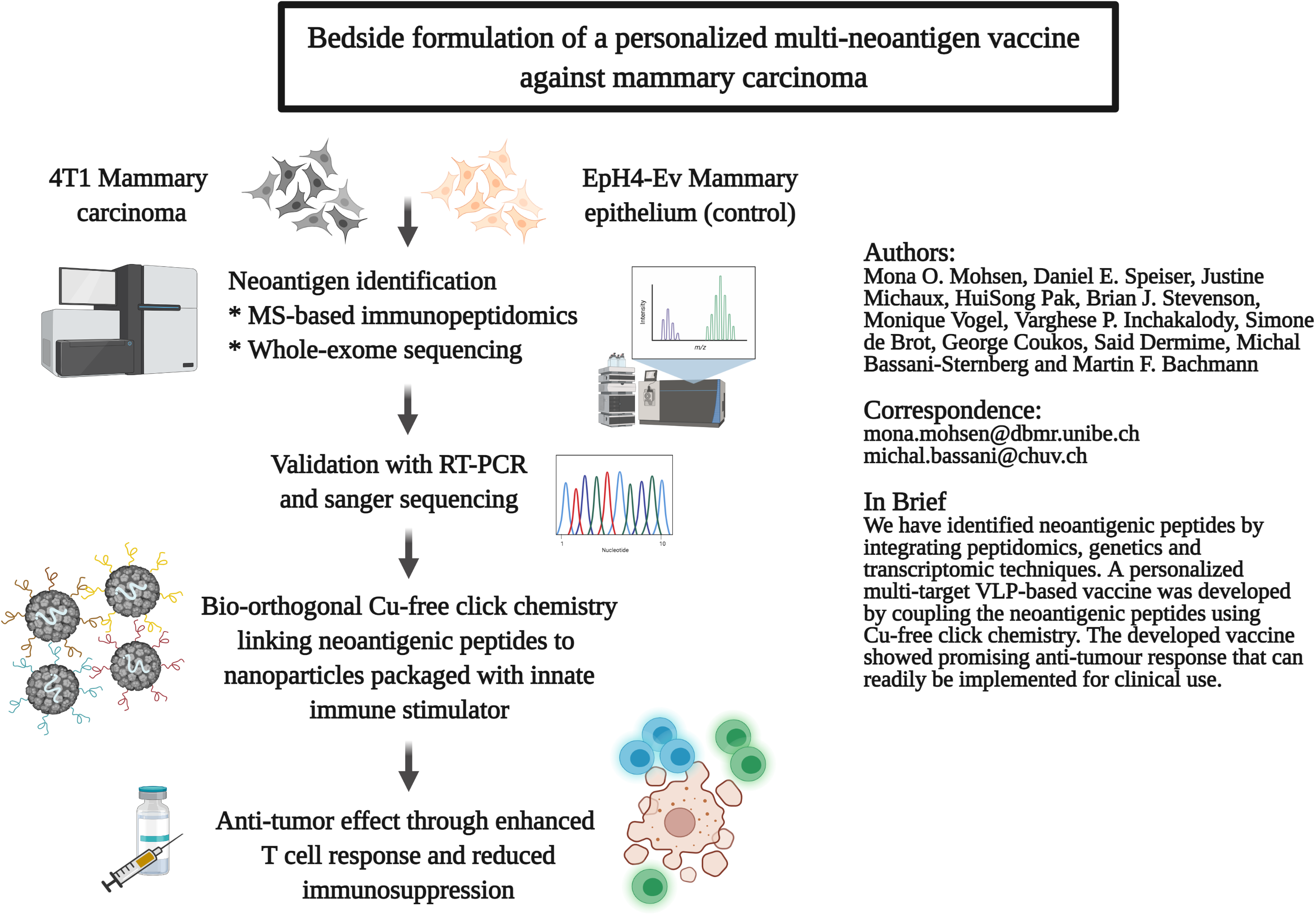

**Background:** Harnessing the immune system to purposely recognize and destroy tumours represents a significant breakthrough in clinical oncology. Nonsynonymous mutations (neoantigenic peptides) were identified as powerful cancer targets. This knowledge can be exploited for further improvements of active immunotherapies, including cancer vaccines as T cells specific for neoantigens are not attenuated by immune tolerance mechanism and do not harm healthy tissues. The current study aimed at developing an optimized multi-target vaccine using short or long neoantigenic peptides utilizing virus-like particles (VLPs) as an efficient vaccine platform.

**Methods:** Here we identified mutations of murine mammary carcinoma cells by integrating mass spectrometry-based immunopeptidomics and whole exome sequencing. Neoantigenic peptides were synthesized and covalently linked to virus-like nanoparticles using a Cu-free click-chemistry method for easy preparation of vaccines against mouse mammary carcinoma.

**Results:** As compared to short peptides, vaccination with long peptides was superior in the generation of neoantigen-specific CD4^+^ and CD8^+^ T cells which readily produced IFN-γ and TNF-α. The resulting anti-tumour effect was associated with favourable immune re-polarization in the tumour microenvironment through reduction of myeloid-derived suppressor cells. Vaccination with long neoantigenic peptides also decreased post-surgical tumour recurrence and metastases, and prolonged mouse survival, despite the tumour’s low mutational burden.

**Conclusion:** Integrating mass spectrometry-based immunopeptidomics and whole exome-sequencing is an efficient technique for identifying neoantigenic peptides. A multi-target VLP-based vaccine shows a promising anti-tumour results in an aggressive murine mammary carcinoma cell line. Future clinical application using this strategy is readily feasible and practical, as click-chemistry coupling of personalized synthetic peptides to the nanoparticles can be done at the bedside directly before injection.

## 1. Introduction

Harnessing the immune system to purposely recognize and destroy tumours represents a significant breakthrough in clinical oncology. Nonsynonymous mutations were identified as powerful cancer targets. This knowledge can be exploited for further improvements of active and passive immunotherapies, including cancer vaccines. T cells specific for neoantigenic peptides are not attenuated by immune tolerance mechanism and do not harm healthy tissues (1). Recent clinical trials have shown that *de novo* immune responses can be induced by vaccination with neoantigens in melanoma and glioblastoma patients (2–5). The optimal peptide length for therapeutic cancer vaccines has been identified in preclinical and clinical studies. Short peptides of ~8-11 amino acids (a.a.) are the typical CD8^+^ T-cell epitopes while long peptides of ~15-32 a.a. contain both CD8^+^ and CD4^+^ T-cell epitopes (6, 7). Neoantigenic epitopes can be predicted based on their sequence, expression, proteasome cleavage, peptide abundance, MHC-binding affinity and other physiochemical and structural features of peptide-MHC complexes. Long peptides are elongated at both ends with natural flanking residues that extend the repertoire of peptides binding to MHC-I to additional peptides binding MHC-II. Anti-tumour immunity induced by cytotoxic CD8^+^ T lymphocytes (CTLs) have been intensely characterized and identified as key players in anti-tumour immunity. Furthermore, CD4^+^ T cells have recently emerged as important contributors in enhancing the immune response against tumours (8, 9).

There exist many different vaccine platforms, based on mRNA, peptides/proteins, dendritic cells (DCs) or recombinant viruses (10–13). In addition, virus like particles (VLPs) are promising, supported by their increasing success as vaccines protecting from various infections (14). Quasicrystalline VLPs are multiprotein complexes with a highly defined repetitive surface geometry, mostly in the shape of an icosahedron (15). Epitope repetitiveness serves as a potent pathogen-associated structural pattern (PASP) efficiently recognized by innate and adaptive immune cells (16). The outer surface of VLPs can be decorated with antigenic peptides. Inside of the VLPs, one can package innate immune stimulators such as toll-like receptor (TLR) ligands for efficient cellular activation (17). VLPs built with bacteriophage Qβ protein (Qβ-VLPs) are highly immunogenic and very versatile for the production of vaccines with antigens and innate immune stimulators. In our earlier studies and clinical trials, we showed that such VLP vaccines induced strong tumour antigen-specific T-cell responses in preclinical models and melanoma patients (18, 19). Our Qβ (G10)-Melan-A vaccine was based on Qβ-VLPs packed with G10, a TLR-9 ligand inducing a potent IFN-α response (20), and decorated with a long peptide derived from the Melan-A^Mart1^ melanoma differentiation antigen. Chemical coupling of peptides to the Qβ-VLPs was performed using the SMPH bifunctional cross-linker. Clinical trials showed that more than half of stage II-IV melanoma patients generated strong tumour antigen-specific and in vivo functional CD8^+^ and CD4^+^ T-cell responses (21). The tumours of some patients showed late loss of Melan-A expression, indicating that targeting of only a single antigen is problematic because of disease progression caused by antigen-loss variants. Indeed, preclinical models confirmed that targeting multiple antigens is superior in generating CD4^+^ and CD8^+^ T-cell responses that can successfully infiltrate tumours, control tumour growth and avoid outgrowth of escape variants (22).

The current study aimed at developing an optimized anti-tumour vaccine using short or long neoantigenic peptides. VLPs were packaged with a TLR-9 ligand and decorated with neoantigenic peptides discovered by peptidomic, genetic and transcriptomic techniques. Vaccines were composed of mixtures of four VLPs each with a different peptide, coupled by click-chemistry permitting easy patient-individual vaccine preparation at the bedside. Our study provides proof of concept of an effective personalized therapeutic cancer vaccine that can readily be implemented for clinical use.

## 2. Materials and methods

### 2.1. Production & purification of Qβ-VLPs

The expression and production of the bacteriophage Qβ-VLPs was carried out as previously described (23–25).

### 2.2. Cell lines and mice

4T1 mammary carcinoma cell line (ATCC® CRL-2539™) and EpH4-Ev wild-type mammary epithelial cell line (ATCC® CRL-3063™) were cultured in DMEM medium supplemented with 10% FBS and 1% penicillin/streptomycin. For EpH4-Ev cell line 6μg Puromycin was added to the culture medium as per ATCC instructions. When cells were 80% confluent, medium was aspirated and cells were washed 3× with 1× PBS to remove excess medium. 1× trypsin was added and flasks were incubated at 37°C for 10min. Cells were collected, resuspended in complete medium and kept on ice until use. Mycoplasma contamination was ruled out using Microsart AMP Mycoplasma Kit. Wild type female BALB/cOlaHsd (8-12 weeks) were purchased from Harlan and kept in the animal facility at the University of Bern. 1×10^6^ cells were inoculated subcutaneously (s.c.) in the flank region under isoflurane anaesthesia. All animal procedures were conducted in accordance with the Swiss Animal Act (455.109.1 – September 2008, 5^th^) of University of Bern.

### 2.3. DNA sequencing and prediction of non-synonymous mutations

DNA was extracted using the commercially available Purelink Genomic DNA mini kit (Invitrogen) as per the manufacturer’s instructions. The purity and concentration of the DNA of both cell lines were measured using nanodrop. Whole exome sequencing (WES) was performed on an Illumina HiSeq 4000 machine using libraries prepared with a SureSelectXT Mouse All Exon capture kit (Agilent Technologies, Santa Clara, CA, USA), to produce 150 paired end reads sufficient for a mean bait coverage of at least 150x. The sequence reads were aligned to the GRCm38.p6 mouse reference assembly using BWA-MEM v0.7.17 (26), and variants were predicted using the Genome Analysis Toolkit best practices protocol (GATK v3.7). A set of reference variants for mouse was derived from data generated by the Wellcome Sanger Institute Mouse Genomes Project (available at ftp://ftp-mouse.sanger.ac.uk/REL-1807-SNPs_Indels/). Somatic mutations (SMs) were defined as variants present only in the 4T1 sample, while single nucleotide polymorphisms (SNPs) were defined as variants present in both the 4T1 and EpH4-Ev samples. The functional effect of the predicted SMs and SNPs was determined by annotating the variants with snpEff v4.3T (27) using the GRCm38.86 and mm10 databases.

### 2.4. Purification of MHC-I binding peptides

Anti–MHC-I monoclonal antibodies were purified from the supernatant of HIB 34.1.2 (a gift from Angel Miguel Garcia-Lora, Hospital Universitario Virgen de las Nieves, Granada, Spain) hybridoma cells cultured in CELLLine CL-1000 flasks (Sigma-Aldrich) using Protein A–Sepharose 4B beads (Invitrogen). Antibodies were cross-linked to Protein A–Sepharose 4B beads at a concentration of 5 mg of antibodies per 1 mL volume of beads. For this purpose, the antibodies were incubated with the Protein A–Sepharose 4B beads for 1 hour at room temperature (RT). Chemical cross-linking was performed by the addition of dimethyl pimelimidate dihydrochloride (Sigma-Aldrich) in 0.2 M sodium borate buffer pH 9 (Sigma-Aldrich) at a final concentration of 20 mM for 30 minutes. The reaction was quenched by incubation with 0.2 M ethanolamine pH 8 (Sigma-Aldrich) for 2 hours. Cross-linked antibodies were kept at 4°C until use. 2.33×10^8^ 4T1cells grown in culture or 8×10^8^ grown in vivo in BALB/c mice per replicate were lysed in phosphate buffered saline containing 0.5% sodium deoxycholate (Sigma-Aldrich), 0.2 mM iodoacetamide (Sigma-Aldrich), 1 mM EDTA, 1:200 Protease Inhibitor Cocktail (Sigma-Aldrich), 1 mM phenylmethylsulfonylfluoride (Roche), and 1% octyl-β-D glucopyranoside (Sigma-Alrich) at 4°C for 1 hour. The lysis buffer was added to the cells at a concentration of 10^8^ cells/mL. Lysates were cleared by centrifugation with a table-top centrifuge (Eppendorf Centrifuge) at 4°C at 20,000 g for 50 minutes. MHC-I molecules were purified sequentially by incubating the cleared lysate first with HIB antibodies cross-linked to protein A-Sepharose 4B beads in affinity columns for 3 hours at 4°C. A ratio of 100 μL of cross-linked beads per 10^8^ cells was applied. The affinity columns were then washed as follows: 2 column volumes of 150 mM sodium chloride (NaCl) in 20 mM Tris-HCl pH 8, 2 column volumes of 400 mM NaCl in 20 mM Tris-HCl pH 8, and again 2 column volumes of 150 mM NaCl in 20 mM Tris-HCl pH 8. Finally, the beads were washed with 1 column volume of 20 mM Tris-HCl pH 8. MHC complexes and the bound peptides were eluted at RT by adding twice a volume of 1% trifluoroacetic acid (TFA) equivalent to or slightly higher than the volume of beads present in the column. Sep-Pak tC18 96-well plates (Waters), preconditioned with 1 mL of 80% acetonitrile (ACN) in 0.1% TFA and then with 2 mL of 0.1% TFA, were used for the purification and concentration of MHC peptides. Elutions containing were loaded in the Sep-Pak tC18 96-well plates and the C18 wells were then washed with 2 mL of 0.1% TFA. The MHC-I peptides were eluted with 500 μL of 28% ACN in 0.1% TFA. The peptide samples were transferred into Eppendorf tubes, dried using vacuum centrifugation (Thermo Fisher Scientific) and stored at −20°C.

### 2.5. Liquid chromatography and mass spectrometry

The LC-MS/MS system consisted of an Easy-nLC 1200 (Thermo Fisher Scientific) hyphenated to a Q Exactive HF-X mass spectrometer (Thermo Fisher Scientific). Peptides were separated on a 450 mm analytical column (8 μm tip, 75 μm inner diameter, PicoTipTMEmitter, New Objective) packed with ReproSil-Pur C18 (1.9 μm particles, 120 Å pore size, Dr. Maisch GmbH). The separation was performed at a flow rate of 250 nL/min by a gradient of 0.1% formic acid (FA) in 80% ACN (solvent B) and 0.1% FA in water (solvent A). MHC-I peptides were analysed by the following gradient: 0–5 min (2-5% B); 5-85 min (5-35% B); 85-100 min (35-60 % B); 100-105 min (60-95% B); 105-110 min (95% B); 110-115 min (95-2% B) and 115-125 min (2% B). For DDA measurements, full MS spectra were acquired in the Orbitrap from m/z = 300-1650 with a resolution of 60,000 (m/z = 200) and an ion accumulation time of 80 ms. The auto gain control (AGC) was set to 3e6 ions. MS/MS spectra were acquired in a data-dependent manner on the 10 most abundant precursor ions with a resolution of 15,000 (m/z = 200), an ion accumulation time of 120 ms and an isolation window of 1.2 m/z. The AGC was set to 2e5 ions. The dynamic exclusion was set to 20 s, and a normalized collision energy (NCE) of 27 was used for fragmentation. No fragmentation was performed for MHC-I peptides with assigned precursor ion charge states of four and above and the peptide match option was disabled.

### 2.6. Identification of neoantigenic peptides from MS immunopeptidomic data

To detect neoantigenic peptides by mass spectrometry, non-synonymous SMs and SNPs were incorporated into the headers of reference protein fasta sequences in a format compatible with MaxQuant 1.5.9.4. as previously described (28). Specifically, mouse GENCODE sequences and the associated gene annotations (vM22; reference assembly version GRCm38) were downloaded from https://www.gencodegenes.org/mouse/releases.html, and all transcripts were scanned for the presence of 4T1-specific variants, using genomic coordinates from the transcript annotations. For each transcript, variants producing non-synonymous changes were identified and the corresponding amino-acid changes were tagged with a unique ID and added to the header of the translated transcript sequence. We searched the immunopeptidomic MS data against the 4T1-specific customized reference database and a file containing 246 frequently observed contaminants with the MaxQuant computational platform version 1.5.9.4. (29). The default settings were used except the following parameters: variants were called from file, enzyme specificity was set to ‘unspecific’, methionine oxidation and Protein N-term acetylation were set as variable modifications and no fixed modification was set, PSM FDR was set to 0.05 with no protein FDR. The initial allowed mass deviation of the precursor ion was set to 6 ppm and the maximum fragment mass deviation was set to 20 ppm. Binding affinity of the peptides to the MHC molecules was predicted by NetMHCpan 4.1 software (30). Binding motif deconvolution of 9 mer MHC-I peptides was performed using the MixMHCp 2.1 (31, 32) with the default settings. Upon completion, deconvoluted motifs were manually assigned as H-2-D^d^ and H-2K^d^.

### 2.7. Verification of identified neoantigens

To verify the expression of the identified neoantigens, RNA from 4T1 mammary carcinoma cell line, 4T1 established tumour and EpH4-Ev wild type mammary epithelial cell line were extracted using NucleoSpin RNA Kit and reverse-transcribed using High-Capacity RNA-to-cDNA Kit as per the manufacturer’s instructions. Three primer pairs were designed of ~24bp each, to amplify the mutated region leading to three amplicons of ~200 bp each. The PCR products were electrophoresed and after gel extraction the amplicons with the expected size were purified using Zymoclean Gel DNA Recovery Kit. Amplicons were then sequenced using Sanger sequencing with the primers used for PCR amplification. The results confirmed the somatic missense mutations in 4T1 mammary carcinoma cell line and the established 4T1 tumours in BALB/cOlaHsd mice, however these mutations were absent from the wild type control mammary epithelial cell line.

### 2.8. Synthesis of neoantigenic peptides (short and long)

The identified neoantigens (NeoAG) in 4T1 mammary carcinoma cell line were synthesized as short (15-16 a.a.) or long (32 a.a.) peptides using their flanking regions as shown in Table 1.

**Table 1.** Synthesized neoantigenic peptides (NeoAG) with flanking regions, additional lysine and an azide group (N3) at the C-terminus.

| Neoantigen (NeoAG) | Class I epitope sequence | Neoantigenic short peptide (NeoAG<sub>S</sub>), 15-16 mers | Neoantigenic long peptide (NeoAG<sub>L</sub>), 32 mers |
| --- | --- | --- | --- |
| NeoAG 1 | KYLDSPKRL | <u>QKQKYLDSPKRL</u>VGLK(N3) | <u>PNTEKITRQKQKYLDSPKRL</u>VGLCGRWNKASK(N3) |
| NeoAG 2 | SPKRLVGL | <u>YLDSPKRLVGL</u>CGRK(N3) | <u>KITRQKQKYLDSPKRLVGL</u>CGRWNKASETLK(N3) |
| NeoAG 3 | KYNAVPTVF | <u>KNLKYNAVPTVF</u>AFQK(N3) | <u>CFSAFGNRKNLKYNAVPTVF</u>AFQNPTEVCPEK(N3) |
| NeoAG 4 | VYREQMNGV | <u>FDSVYREQMNGV</u>QRFK(N3) | <u>FTVAQSEAFDSVYREQMNGV</u>QRFPWDTSEEDK(N3) |

An additional lysine (K) and an azide group (N3) were added at the C-terminus of the synthesized peptides to facilitate Cu-free click chemistry coupling. Neoantigenic short and long peptides are referred to as NeoAG**_S_** and NeoAG**_L_**, respectively. Neoantigenic MHC-I peptides (8 or 9 a.a. in blue), non-synonymous somatic mutations (highlighted in grey), flanking regions (in blue), additional lysine (K) (in red) and the azide group (N3) (in black).

### 2.9. Production of Qβ(1668)-NeoAG_S_ and Qβ(1668)-NeoAG_L_ vaccines

The naturally packaged ssRNA in Qβ-VLPs was digested as following: Qβ(ssRNA)-VLPs were washed 3x with 20mM HEPES (Sigma-Aldrich) using Amicon ultra 0.5ml tubes 100kDa MWCO. RNAse A (Merck) was used to digest the ssRNA (1.2mg/ml for each 3mg/ml Qβ-VLPs) and incubated at 37°C for 3h in a shaker 400rpm. The digestion of the ssRNA was confirmed by 1% agarose gel stained with SYBR Safe dye for 30min at 90V. Qβ-VLPs were then re-packaged with B-type 1668 CpGs (5″-TCC ATG ACG TTC CTG ATG CT-3″) with phosphorothioate backbone purchased from (InvitroGen). The re-packaging was confirmed by 1% agarose gel stained with SYBR Safe dye for 30min at 90V. The presence of Qβ(1668)-VLPs protein was detected by staining the 1% agarose gel with Coomassie blue stain. Qβ(1668)-VLPs were then derivatized for 30min at RT using dibenzocyclooctyne NHS ester (DBCO cross-linker) (Sigma-Aldrich) in a 5-fold molar excess. Amicon ultra 0.5ml tubes 100kDa MWCO were used to remove excess DBCO cross-linker. The NeoAG_S_ or NeoAG_L_ peptides were synthesized by (Pepscan BRESTO) with the addition of lysine (K) and (N3) azide group at the C-terminus to facilitate their coupling to the derivatized Qβ(1668)-VLPs. The synthesized peptides were reconstituted in DMSO and added in a 4-fold molar excess over Qβ(1668)-VLPs monomer. The vaccine was incubated 1h at RT and Amicon ultra 0.5ml tubes 100kDa MWCO were used to remove excess peptide. The coupling of the NeoAG_S_ or NeoAG_L_ to Qβ(1668)-VLPs was done separately for each neoantigenic peptide. The efficiency of the coupling was checked with SDS-PAGE (BIO-RAD).

### 2.10. Vaccination of mice

BALB/cOlaHsd mice were vaccinated after inoculation of 4T1 mammary carcinoma cell line as described in 2.2. Designed groups, vaccination doses and regimen were scheduled as shown in the results section. Briefly, we injected 1X10^6^ 4T1 s.c. in the flank as previous studies have shown that this method possesses an increased localization of the primary tumour which makes it easier to be resected in surgery. Injecting 4T1 in mammary fat or orthotopically showed enhanced invasive growth pattern. The s.c. inoculation method of 4T1 results in metastases as per our primary data and as shown by other studies (33). Vaccination started 3 days after cell line inoculation as summarized in Table 2. Mice were followed every 2 days to assess tumour volume and general health score. Three doses (Prime-Boost-Boost) of the vaccine (40μg each dose) were administered over 17 days (please refer to results 3.3). Tumours in the control group reached the ethically maximal tolerated size of 1 cm^3^ by day 17. For the experiment in 3.6, primary tumours were resected on day 14 from all groups and 2 vaccine doses were administered weekly. Mice weight was followed routinely and were euthanised when loosing 10-15% of their original weight. Lungs were collected when mice were euthanized and lung weight was measured.

**Table 2.**
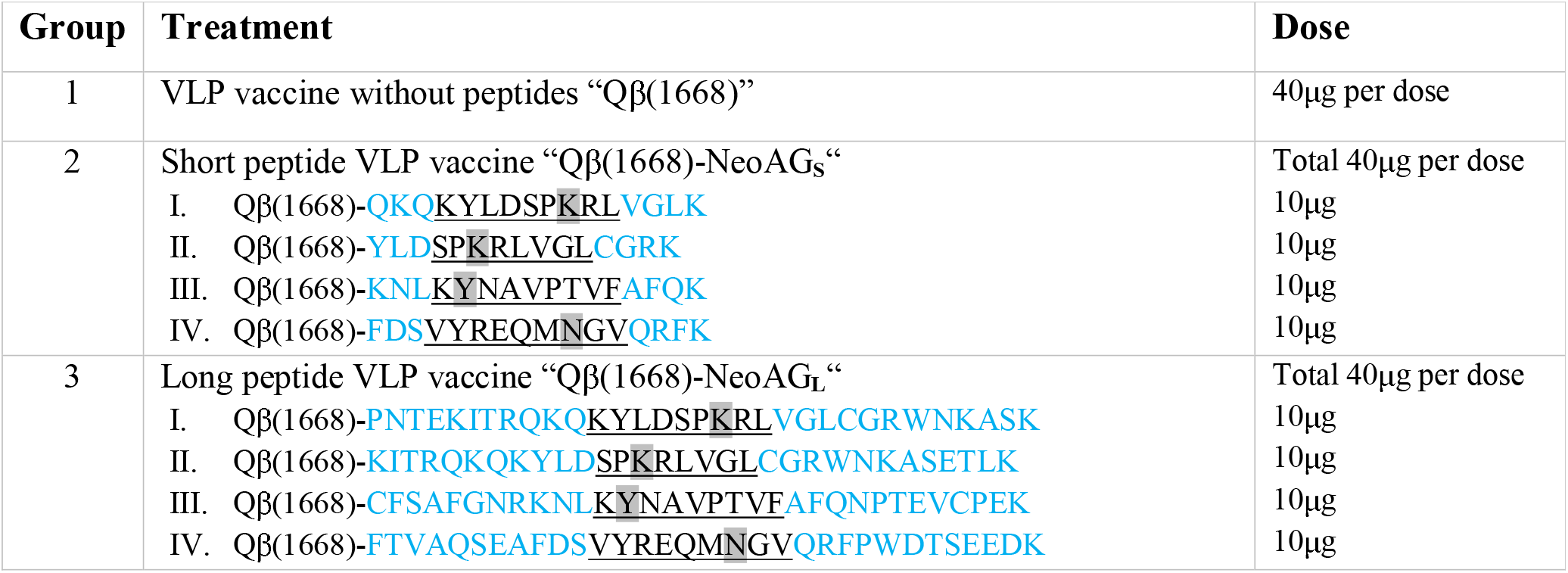
Treatment groups and vaccine doses.

### 2.11. Analysis of blood cells

Blood was collected from mice’ tail vein on day 16 after inoculation of 4T1 mammary carcinoma. 150μl blood was collected in 500μl 1x PBS containing heparin and kept on ice. Cells were centrifuged at 1200rpm for 5min and supernatant was aspirated. RBCs were lysed using 500μl ACK buffer (Sigma-Aldrich) on ice for 2-3 min. White blood cells (WBCs) were collected after 5min centrifugation at 1200rpm. Supernatant was aspirated and cells were resuspended with 1x PBS containing 0.1% BSA and again centrifuged. Pelleted cells were labelled with live/dead stain, anti-mouse CD16/CD32 (mouse BD Fc block) mAb clone 2.4G2 (BD Bioscience) for 10min in the dark, centrifuged as described above and stained with PE anti-mouse CD8α mAb clone 53-6.7 (BD Bioscience), PE/Cyanine7 anti-mouse CD4 mAb clone RM4-5 (Biolegend). Samples were read by FACSCaliber and analysis was done using GraphPad Prism version 8.4.2 (464).

### 2.12. Analysis of tumours

Tumours were measured every 3 days using a calliper using the formula V=(WxWxL)/2, V=final tumour volume in mm^3^, L=tumour length and W=tumour width. On day 17 mice were euthanised as tumours in the control group reached the size of 1cm^3^. Tumours were collected in DMEM medium containing 10%FBS and 1% Penicillin/Streptomycin on ice. Tumour infiltrating lymphocytes (TILs) were collected as following: tumours were dissected into pieces and smashed using 70μM cell strainer, cells were washed during the process using DMEM medium containing 10%FBS and 1% Penicillin/Streptomycin in falcon tubes 50ml. Collected cells were added to 15ml tubes containing 2ml of 35% Percoll slowly. The tubes were centrifuged at 1800rpm for 25min at RT to isolate TILs. TILs were then resuspended in 200μl PBS (0.1% BSA) and 100μl was transferred to 96 well v-bottom plates and centrifuged at 1200rpm for 5min. Supernatant was discarded, and RBCs were lysed using ACK 500μl ACK buffer (Sigma-Aldrich) on ice for 2-3 min. TILs were stained with live/dead, anti-mouse CD16/CD32 (mouse BD Fc block) mAb clone 2.4G2 (BD Bioscience) for 10min in the dark for 10min, centrifuged as described above and stained with PE anti-mouse CD8α mAb clone 53-6.7 (BD Bioscience) and PE/Cyanine7 anti-mouse CD4 mAb clone RM4-5 (Biolegend). In the second experiment, TILs were also stained with APC anti-mouse Ly6G mAb clone 1A8 (BioLegend), FITC anti-mouse Ly6C clone HK1.4 (BioLegend) and APC/Cyanine7 anti-mouse CD11b clone M/170 (BioLegend). Plates were centrifuged at 1200rpm for 5min, supernatant was discarded, TILs were resuspended in PBS (0.1% BSA) and added to 5ml round-bottom tubes with cell strainer to remove excess tumour debris. Samples were read by FACSCaliber and analysis was done using GraphPad Prism version 8.4.2 (464).

### 2.13. Intra cellular cytokine staining (ICS)

100μl of the TILs described in 2.12 were transferred to sterile 96 well flat-bottom plates. TILs were incubated with mouse IL-2 (mIL2-Ref: 11271164001-MERCK) 100U/ml in DMEM medium containing 10% FBS and 1% Penicillin/Streptomycin at 37°C for 2 days. TILs were washed 3x with DMEM medium containing 10% FBS and 1% Penicillin/Streptomycin and a stimulation cocktail was added 1μg/ml of each NeoAG with Brefeldin and Monensin (1:1000) at 37°C for 6h. TILs were washed 3x with DMEM medium to remove the stimulation cocktail and then transferred to 96 well v-bottom plates for staining. TILs were stained with live/dead, anti-mouse CD16/CD32 (mouse BD Fc block) mAb clone 2.4G2 (BD Bioscience) for 10min in the dark, centrifuged as described above and stained with PE anti-mouse CD8α mAb clone 53-6.7 (BD Bioscience) and PE/Cyanine7 anti-mouse CD4 mAb clone RM4-5 (Biolegend). The plates were centrifuged at 1200rpm for 5min, supernatant was discarded and TILs were fixed using 100μl of the fixation buffer (BD Cytofix) at 4°C for 15min. The plates were again centrifuged at 1200rpm for 5min, supernatant was discarded, and TILs were washed with 100μl of 1x diluted permeabilization wash buffer (BioLegend) and centrifuged immediately at 1200rpm for 5min, supernatant was discarded. TILs were then stained with APC anti-mouse IFN-γ mAb clone XMG1.2 (MERCK) and PerCP-Cyanine5.5 anti-mouse TNF-α mAb clone MP6-XT22 (BioLegend). Plates were centrifuged at 1200rpm for 5min, supernatant was discarded, TILs were resuspended in PBS (0.1% BSA) and added to 5ml round-bottom tubes with cell strainer to remove excess tumour debris. Samples were read by FACSCaliber and analysis was done using GraphPad Prism version 8.4.2 (464).

### 2.14. Immunohistochemistry

Formalin-fixed murine mammary carcinomas were assessed by immunohistochemistry (IHC) staining for CD4^+^ and CD8^+^ cells using CD4 (clone 4SM95, rat, Thermo F. Scientific, 14-9766) and CD8 (rat, Dianova, DIA-808). For each tumour, one full cross section was examined by a board-certified veterinary pathologist (SdB). The quantitative evaluation of peritumoral and intratumoral CD4 and CD8 positive cells was done digitally using the software QuPath 0.2.3. The number of CD4^+^ and CD8^+^ cells was calculated per mm (tumour periphery) and per mm2 (intratumoral area).

### 2.12. Statistics

Data were presented as mean ± SEM. Comparisons between groups was performed by Student’s unpaired *t* test (2-tailed). Area under curve (AUC) was used to measure tumour growth curves. Survival rate was analysed by Log-rank (Mantel-Cox) test. P values \*\*\*\**P <* 0.0001; \*\*\**P <* 0.001; \*\**P <* 0.01; \**P <* 0.05.

## 3. Results

### 3.1. Neoantigen identification

The current study aimed at providing a proof of concept for the generation of a personalized virus-like particle (VLP)-based vaccine by incorporating tumour-specific neoantigens of mammary carcinoma. We have chosen the aggressive, highly metastatic and low mutational burden mammary carcinoma cell line (4T1) (11). The wild type mammary epithelial cell line, EpH4-Ev, was selected as germline reference cells. Both cell lines originate from BALB/cOlaHsd mice. Neoantigen identification was performed by whole exome sequencing combined with immunopeptidomics (Fig. 1A). MHC-I binding peptides were eluted from 4T1 cells and were characterized by LC-MSMS. A 4T1-specific reference database that includes non-synonymous somatic mutations was generated to match the MSMS data. In total, 2714 and 5077MHC-I peptides were identified in the 4T1 mammary carcinoma cells grown *in vitro* and *in vivo*, respectively (Supplementary Data 1). The peptides recapitulated the expected length distribution of MHC-I ligands, meaning the majority were 9 mers, and the 9 mer peptides were clustered into two groups, revealing the expected binding motifs of H-2D^d^ and H-2-K^d^ (Fig. 1B-D). Despite the low mutational burden of 4T1, we identified 4 neoantigens by MSMS (Fig. 1E-H). The non-synonymous somatic mutations were identified in Vrk3 (E391K), Thap3 (H75Y) and Sult2b1 (H299N) as shown in Figure. 1, and Supplementary Table 1.

**Figure 1.**
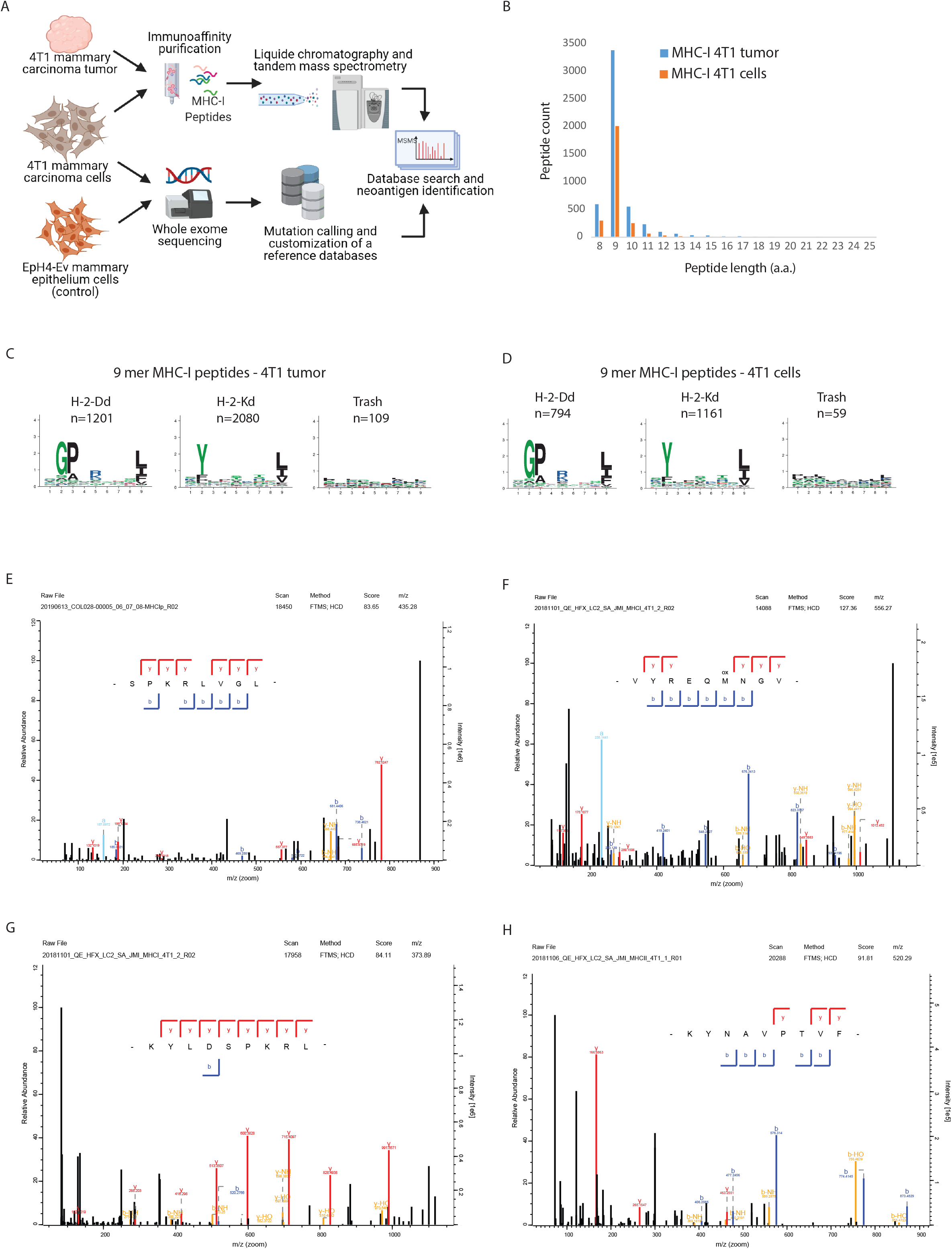
Neoantigen identification. *A*, Schematic representation of our approach for neoantigen identification. *B*, Length distribution of MHC-I peptides identified by immunopeptidomics from 4T1 cells grown in vivo or in vitro. *C*, Clustering of 9 mer MHC-I peptides identified in 4T1 cells grown in vivo and *D*, in 4T1 cells grown in vitro. *E-H*, annotated MSMS spectra of the neoantigens identified by MaxQuant in 4T1 cells. MSMS spectra were exported from the MaxQuant viewer.

### 3.2. Verification of neoantigen sequence and expression

We performed RT-PCR to verify the sequence and expression of the neoantigens. RNA was extracted from the 4T1 mammary carcinoma cell line, from 4T1 tumour *in vivo* and from the EpH4-Ev wild type mammary epithelial cell line as control. The results confirmed presence and expression of the non-synonymous somatic mutations in 4T1 mammary carcinoma cell line as well as in 4T1 tumour *in vivo* in BALB/c mice. These mutations were absent in EpH4-Ev cells (Supplementary Fig. 1).

### 3.3. Efficient coupling of synthetic NeoAG peptides by Cu-free click chemistry to VLPs

Qβ(ssRNA)-VLPs were expressed and purified as described previously (23, 24). The integrity of Qβ(ssRNA)-VLPs was confirmed by electron microscopy (Fig. 2A). In a next step, Qβ(ssRNA)-VLPs were treated with RNaseA to digest the RNA that was naturally packaged during protein expression in *E. coli* (17, 34). The empty particles were then re-packaged with the B-type CpGs 1668, a Toll-like receptor (TLR-9) ligand and potent activator of DCs required for effective DC maturation and cross-priming. The re-packaging is based on direct diffusion of CpG into Qβ-VLPs via its natural pores followed by binding to arginine repeats inside the capsid which in the natural virus are binding the viral genome (35). The re-packaging process of Qβ-VLPs with the CpGs was confirmed in a 1% agarose gel stained with SybrSAFE (Fig. 2B-top), and presence of the VLP’s capsid protein (CP) was assessed by staining the agarose gel with Coomassie-blue (Fig. 2B-bottom). In two previous and independent studies we have shown that the bio-orthogonal Cu-free click chemistry is an efficient, safe (non-toxic) and fast method to chemically couple peptides of ~14 a.a. to Qβ-VLPs (22) or to the plant-derived virus CuMV_TT_-VLPs (36). For several reasons, including ease of manufacturing and safety, this method is preferable over using the heterobifunctional cross-linker SMPH when considering the development of a personalized cancer vaccine (see also discussion). Short (15-16 a.a.) or long (32 a.a.) neoantigenic peptides were synthesized with an additional lysine (K) and an azide group (N3) at the C-terminus (Table 1) and coupled to Qβ(1668)-VLPs (Table 2). The coupling efficiency was confirmed by SDS-PAGE followed by densitometric analysis. SDS-PAGE showed Qβ-VLP monomers (~14kDa) as well as additional bands, indicating effective coupling of the synthesized neoantigens (NeoAG) to Qβ(1668)-VLPs (Fig. 2C). The densitometric analysis of Qβ(1668) alone, Qβ(1668)-NeoAG**_S_** and Qβ(1668)-NeoAG**_L_** confirmed that the coupling was efficient (data not shown). Each NeoAG peptide (short or long) was coupled to Qβ(1668)-VLPs separately. To avoid possible outgrowth of any tumour escape variants, we prepared the final vaccines as multi-target neoantigen vaccines (Fig. 2D). The vaccine with the short peptides was termed Qβ(1668)-NeoAG_S_ and the vaccine with the long peptides Qβ(1668)-NeoAG_L_, each containing one of the neoantigenic peptides with the mutations identified in Vrk3 (E391K), Thap3 (H75Y) or Sult2b1 (H299N). The mutation in Vrk3 (E391K) was retrieved in two different MHC-I neoepitopes. The sequences and treatment doses are shown in Table 2.

**Figure 2.**
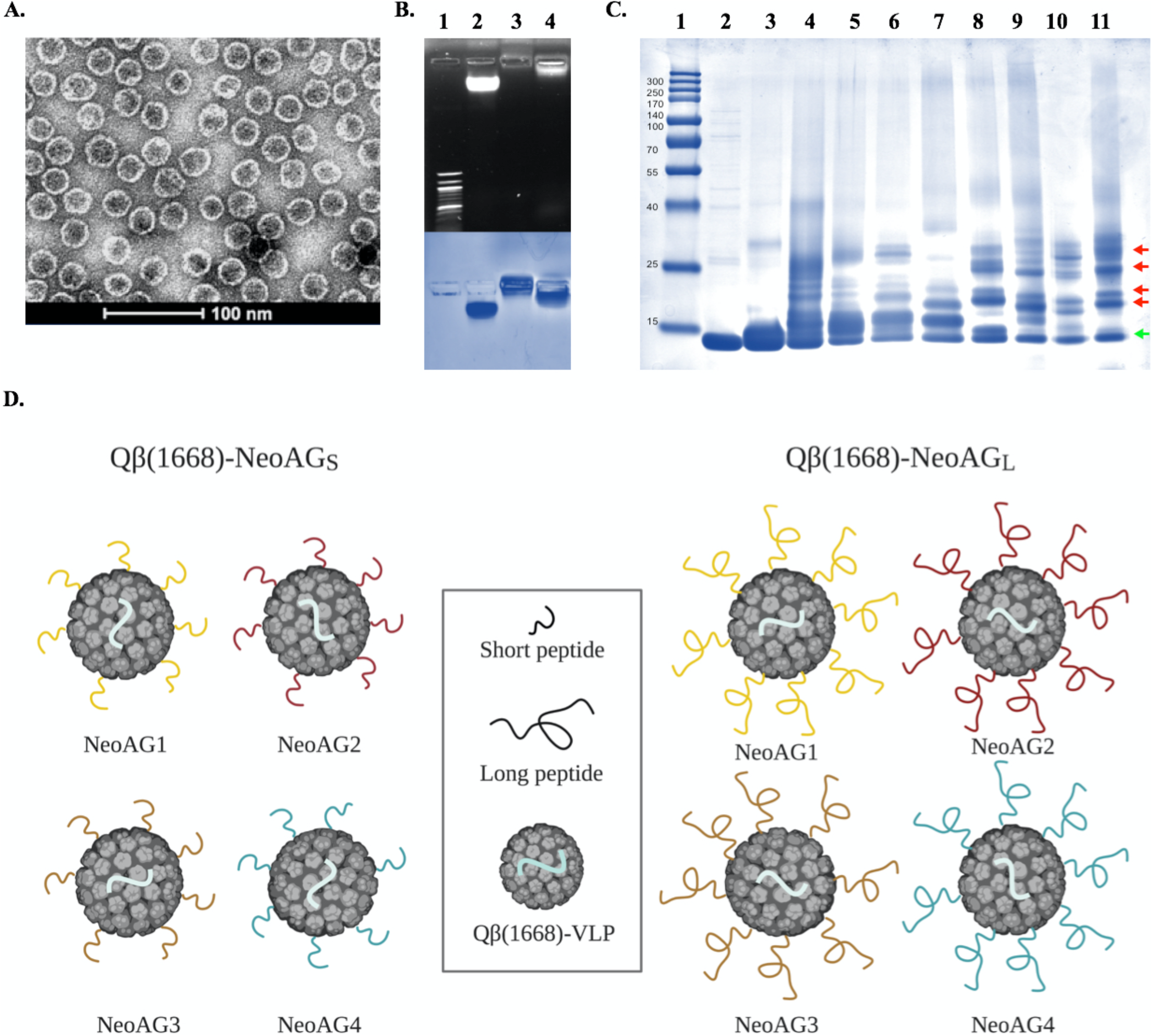
Efficient coupling of synthetic NeoAG peptides by Cu-free click chemistry to VLPs. *A*, Electron microscopy imaging of the Qβ(ssRNA)-VLPs used for vaccine development. *B*, (top) 1% agarose gel stained with SybrSafe, Lane 1: DNA marker, Lane 2: 10μg of Qβ(ssRNA)-VLPs, Lane 3: 10μg of Qβ(empty)-VLPs digested with RNase A, Lane 4: 10μg of Qβ(1668)-VLPs re-packaged with B-type CpGs, (Bottom) Coomassie Blue staining of the agarose gel to confirm integrity of the VLPs, similar lanes as indicated on top. *C*, SDS-PAGE stained with Coomassie Blue, Lane 1: protein ladder, Lane 2: 6μg of Qβ(1668)-VLPs, Lane 3: 6μg of Qβ(1668)-VLPs derivatised with Cu-free click chemistry cross-linker (DBCO), Lanes 4-7: 6μg of Qβ(1668)-VLPs derivatised with Cu-free click chemistry cross-linker (DBCO) and coupled to NeoAG1 or NeoAG2 or NeoAG3 or NeoAG4 (15-16 a.a. short peptides), Lanes 8-11 6ug of Qβ(1668)-VLPs derivatised with Cu-free click chemistry cross-linker (DBCO) and coupled to NeoAG1 or NeoAG2 or NeoAG3 or NeoAG4 (32 a.a. long peptides). The green arrow points to Qβ(1668)-VLP monomers, red arrows point to a peptide bound to Qβ monomer. *D*, Cartoon illustrating the multi-target vaccines: Qβ(1668)-NeoAG_S_ (left) and Qβ(1668)-NeoAG_L_ (right).

### 3.4. Superior *in vivo* efficacy by the long as opposed to the short peptide vaccine

We compared the *in vivo* anti-tumour efficacy of the short and long peptide vaccines. To this end, the aggressive 4T1 mammary carcinoma cell line was inoculated s.c. in Balb/c mice as described in 2.2. On days 3, 7 and 12, s.c. vaccinations were done with Qβ(1668)-NeoAG_S_, Qβ(1668)-NeoAG_L_, or Qβ(1668) as control without peptides (Fig. 3A). Vaccination with the Qβ(1668)-NeoAG_S_ significantly (*p*. 0.0133) hindered tumour progression as compared to the Qβ(1668) control. Interestingly, the long peptide vaccine Qβ(1668)-NeoAG_L_ was even more potent than the short peptide vaccine (*p.* 0.0091) and largely superior to the control group (*p*. 0.0001) (Fig. 3B-D). It is well known that infiltration of CD8^+^ T cells into the tumour correlates with better prognosis which serves as an essential piece of evidence for effective immune responses in melanoma (37, 38) as well as breast cancer (39). Furthermore, it has been suggested that the majority of immunogenic neoantigens in personalized cancer vaccines activate CD4^+^ T cells (40). Accordingly, we determined the density of CD4^+^ and CD8^+^ T cells in each tumour. No significant differences were found for CD8^+^ (*p*. 0.9759) or CD4^+^ (*p*. 0.847) T-cell infiltration between the control group and the group treated with the short peptide vaccine. Yet, a considerably increased CD4^+^ T-cell infiltration was observed after long peptide vaccination in comparison to the control group or the short peptide vaccinated group (Fig. 3E,G). The infiltration of CD8^+^ T cells was also increased after long peptide vaccination (Fig. 3F,H). To study the systemic effects of the vaccinations we characterized CD4^+^ and CD8^+^ T cells in peripheral blood fifteen days post 4T1 inoculation. We found no significant differences between the groups (data not shown). We also assessed myeloid-derived suppressor cells (MDSCs) characterized by CD11b^Hi^ Ly6C^Hi^ or CD11b^Hi^ Ly6G^Hi^ (Fig. I,J). Interestingly, the percentage of CD11b^Hi^ Ly6C^Hi^ was significantly decreased in the group treated with Qβ(1668)-NeoAG_S_ when compared to the control group (*p*. 0.0132). The long peptide vaccine Qβ(1668)-NeoAG_L_ was even more efficient in diminishing this population in comparison to the control group (*p*. <0.0001) or the group treated with the short peptide vaccine (*p*. 0.003) (Fig. 3I,L).

**Figure 3.**
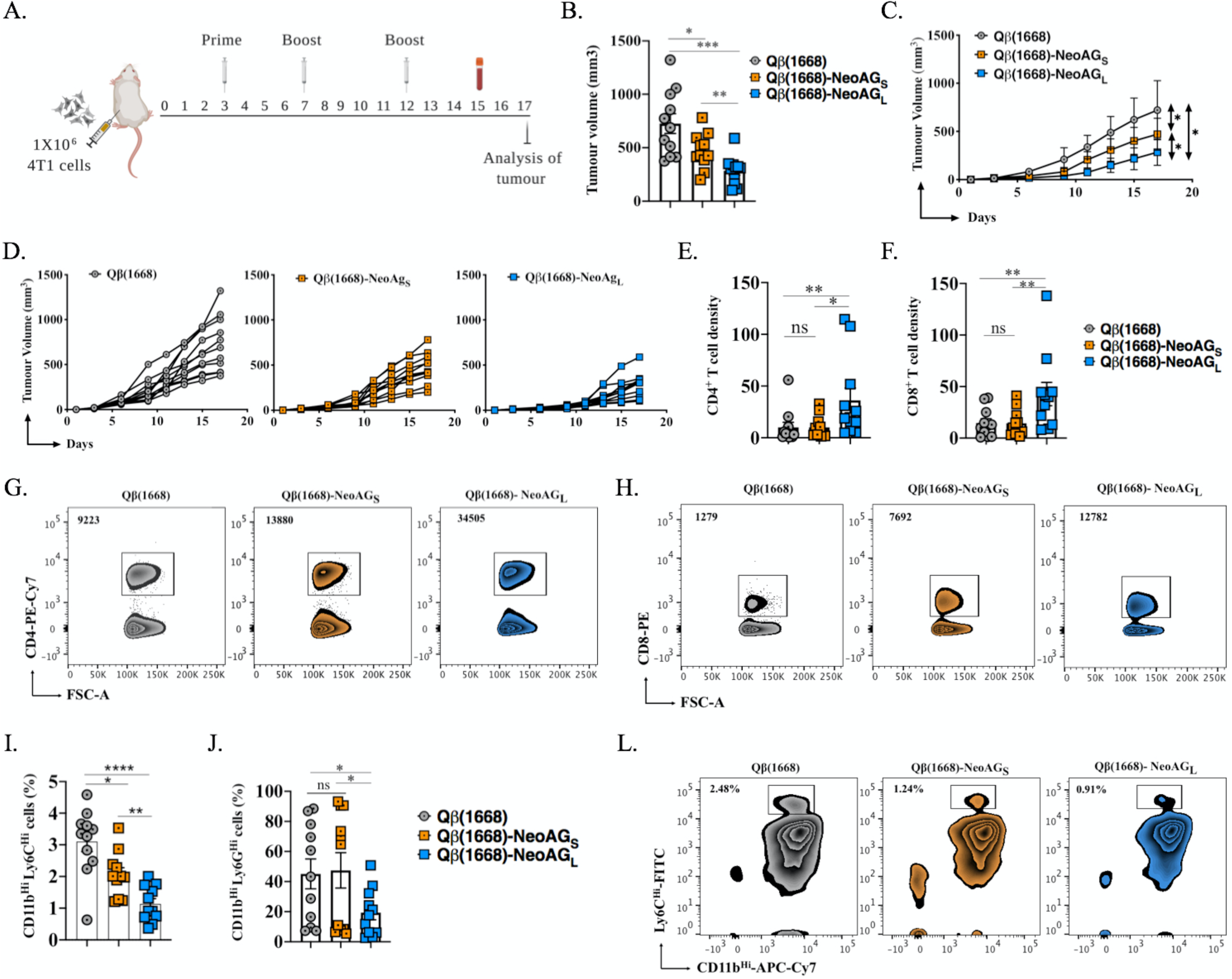
Superior *in vivo* efficacy by the long as opposed to the short peptide vaccine. *A*, Treatment regimen using a prime and 2 boost vaccinations as summarized in Table 2. *B*, Tumour volume in mm^3^ (means ± SEM) measured on day 15 post 4T1 inoculation. Each dot represents an individual tumour. *C*, Tumour growth curves of the treated groups, AUC was measured and statistical analysis was done by Student’s *t*-test (2-tailed). D, Individual tumour growth curves of the treated groups. *E,F*, Density of CD4^+^ or CD8^+^ T cells (means ± SEM) in tumours as determined by flow cytometry. The densities were determined by dividing the total number of CD4^+^ or CD8^+^ T cells in each tumour by the tumour volume in(mm3), pre-gated on tumour infiltrating lymphocytes (TILs). *G,H*, Representative FACS plots showing the percentage of CD4^+^ or CD8^+^ T cells in tumours, pregated on TILs. *I,J*, Percentage of CD11b^Hi^ Ly6C^Hi^ or CD11b^Hi^ Ly6G^Hi^ cells in peripheral blood 15 days post 4T1 inoculation. *K,L*, Representative FACS plots showing the percentage of CD11b^Hi^ Ly6C^Hi^ or CD11b^Hi^ Ly6G^Hi^ in peripheral blood 15 days post 4T1 inoculation, pre-gated on monocytes. Statistical analysis by Student’s *t*-test (2-tailed), (n=11), one representative of 2 similar experiments is shown. *P* values \*\*\*\**P <* 0.0001; \*\*\**P <* 0.001; \*\**P <* 0.01; \**P <* 0.05.

### 3.5. In the tumour microenvironment, the long peptide vaccine reduced MDSCs and increased IFN-γ and TNF-α production by T cells

Based on its superiority we focused on the long peptide vaccine Qβ(1668)-NeoAG_L_. In order to characterize the intratumoural immune response induced by vaccination, we compared tumours from control mice with those after vaccination with the long peptide vaccine (Qβ(1668)-NeoAG_L_). The results confirmed that long peptide vaccination can significantly hinder 4T1 tumour growth (*p*. 0.0022) when compared to the control group treated with Qβ(1668) (Fig. 4A). T-cell density was assessed in each tumour by flow cytometry. We found significantly enhanced intratumoural infiltration of CD4^+^ and CD8^+^ T cells (*p*. 0.0035 and 0.0005 respectively) (Fig. 4B,C), inversely correlating with tumour volume (*p*. 0.0089 and 0.0079, respectively) (Fig. 4D,E). Furthermore, we determined the production of IFN-γ and TNF-α by CD4^+^ and CD8^+^ T cells after stimulation of T cells with a cocktail of the long peptides. The results showed significant increase of IFN-γ or TNF-α in specific CD4^+^ T cells (*p*. 0.0022 and 0.0303, respectively) (Fig. 4F,G) after long peptide vaccination compared to controls. This was also the case for the specific polyfunctional CD4^+^ T cells that simultaneously produced both cytokines (*p*. 0.0022) (Fig. 4H,L). Production of IFN-γ by specific CD8^+^ T cells was also significantly increased (*p*. 0.0022) (Fig. 4I). Despite a trend, the difference for TNF-α production was not significant, likely related to the high heterogeneity (Fig. 4J). When assessing specific CD8^+^ T cells producing both cytokines, a significant increase was found after long peptide vaccination as compared to control mice (*p*. 0.0022) (Fig. 4K,M). Finally, we also studied the effect of vaccination on intratumoural monocytic and granulocytic myeloid cells. Interestingly, both the CD11b^Hi^ Ly6C^Hi^ and the CD11b^Hi^ Ly6G^Hi^ cells were significantly decreased after long peptide vaccination (Fig. 4N,O).

**Figure 4.**
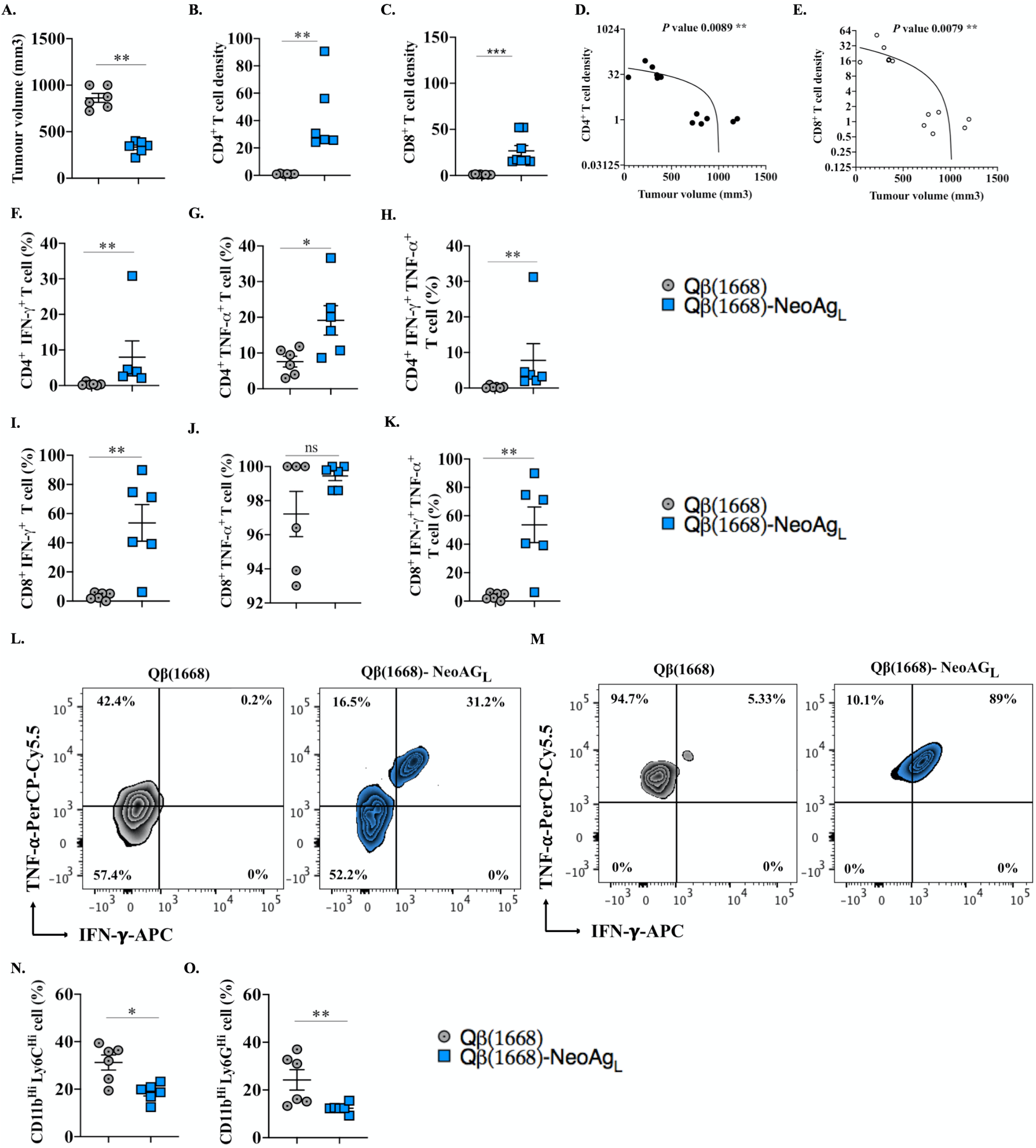
In the tumour microenvironment, the long peptide vaccine reduced MDSCs and increased IFN-γ and TNF-α production by T cells. *A*, Tumour volume in mm^3^ (means ± SEM) measured on day 17 post 4T1 inoculation of control mice treated with Qβ(1668) or mice vaccinated with Qβ(1668)-NeoAG**_L_**. Each dot represents an individual tumour. *B,C*, CD4^+^ or CD8^+^ T cells density (means ± SEM) in tumours as determined by flow cytometry and dividing the total number of CD4^+^ or CD8^+^ T cells in each tumour by the tumour volume in (mm3). *D,E*, Correlations between CD4^+^ or CD8^+^ T cells density in each tumour and its corresponding tumour volume in (mm3). *F,G,H*, Percentage of CD4^+^ IFN-γ^+^, CD4^+^ TNF-α^+^ and dual IFN-γ^+^, TNF-α^+^ CD4^+^ T cells (means ± SEM) in tumours. *I, J, K*, Percentage of CD8^+^ IFN-γ^+^, CD8^+^ TNF-α^+^ and dual IFN-γ^+^, TNF-α^+^ CD8^+^ T cells (means ± SEM) in tumours. *L, M*, Representative FACS plots showing the percentage of dual IFN-γ^+^, TNF-α^+^ CD4^+^ and CD8^+^ T cells in tumours. *N,O*, Percentage of CD11b^Hi^ Ly6C^Hi^ cells and CD11b^Hi^ Ly6G^Hi^ cells (means ± SEM) in tumours. Statistical analysis by Student’s *t*-test. (n=6), one representative of two similar experiments is shown for each data set. P values \*\*\*\**P <* 0.0001; \*\*\**P <* 0.001; \*\**P <* 0.01; \**P <* 0.05.

### 3.6. Long peptide vaccination decreases post-surgical tumour recurrence and lung metastases, and prolongs mouse survival

The observed efficacy of vaccination with Qβ(1668)-NeoAG_L_ *in vivo* promoted us to address whether the vaccine may be useful in a clinically relevant perspective, namely to prevent tumour progression after surgical resection of the primary tumour. To this end, 1 million 4T1 cells were inoculated in the left flank of the mice. Treatment with Qβ(1668) as a control or with the vaccine Qβ(1668)-NeoAG_L_ was initiated 3 days later. Mice received 3 doses over 13 days and primary tumours were surgically resected on day 14, followed by two treatment doses weekly (Fig. 5A). Qβ(1668)-NeoAG_L_ hindered 4T1 primary tumour progression as shown in (Fig. 5B,C) confirming our findings. By immunohistochemistry we investigated CD4^+^ and CD8^+^ T-cell infiltration into the primary tumours (Fig. 5D). We found a significant increase (*p*. 0.0229) in the numbers of peritumoural CD4^+^ T cells in the group vaccinated with Qβ(1668)-NeoAG_L_ in comparison to the control group (Fig. 4E). The numbers of intratumoural CD4^+^ T cells was heterogenous and showed a trend but no significant increase in the group vaccinated with Qβ(1668)-NeoAG_L_ (Fig. 5F and Supplementary Fig. 2). After surgical resection of primary tumours, the weight of mice in the control group remained stable for 10 days (until day 24 of primary tumour inoculation), but started to drop sharply thereafter when all the mice of the control group had to be sacrificed at days 39-40 after primary tumour inoculation. The mice vaccinated with Qβ(1668)-NeoAG_L_ showed much less weight loss and remained stable after day 40 (for the surviving mice) (Fig. 5G). However, the weight loss in some of the immunized mice required that 20% of the mice had to be euthanised on day 42 and 20% on day 48, whereas the remaining 60% of mice continued completely healthy and had stable weight until the end of the experiment on day 66 (Fig. 5H). We measured the weight of the lungs of euthanised mice to quantify lung metastases. The results reflect extensive 4T1 lung metastases in the control group, whereas long peptide vaccination resulted in significantly less lung metastases in the mice that were euthanised due to weight loss, and no lung metastases in the remaining 60% of the mice at the end of the experiment (*p*. 0.0159) (Fig. 5I,J). Analysing the volume of the primary tumour and lung weight revealed a significant correlation (*p*. 0.0268) (Fig. 5K).

**Figure 5.**
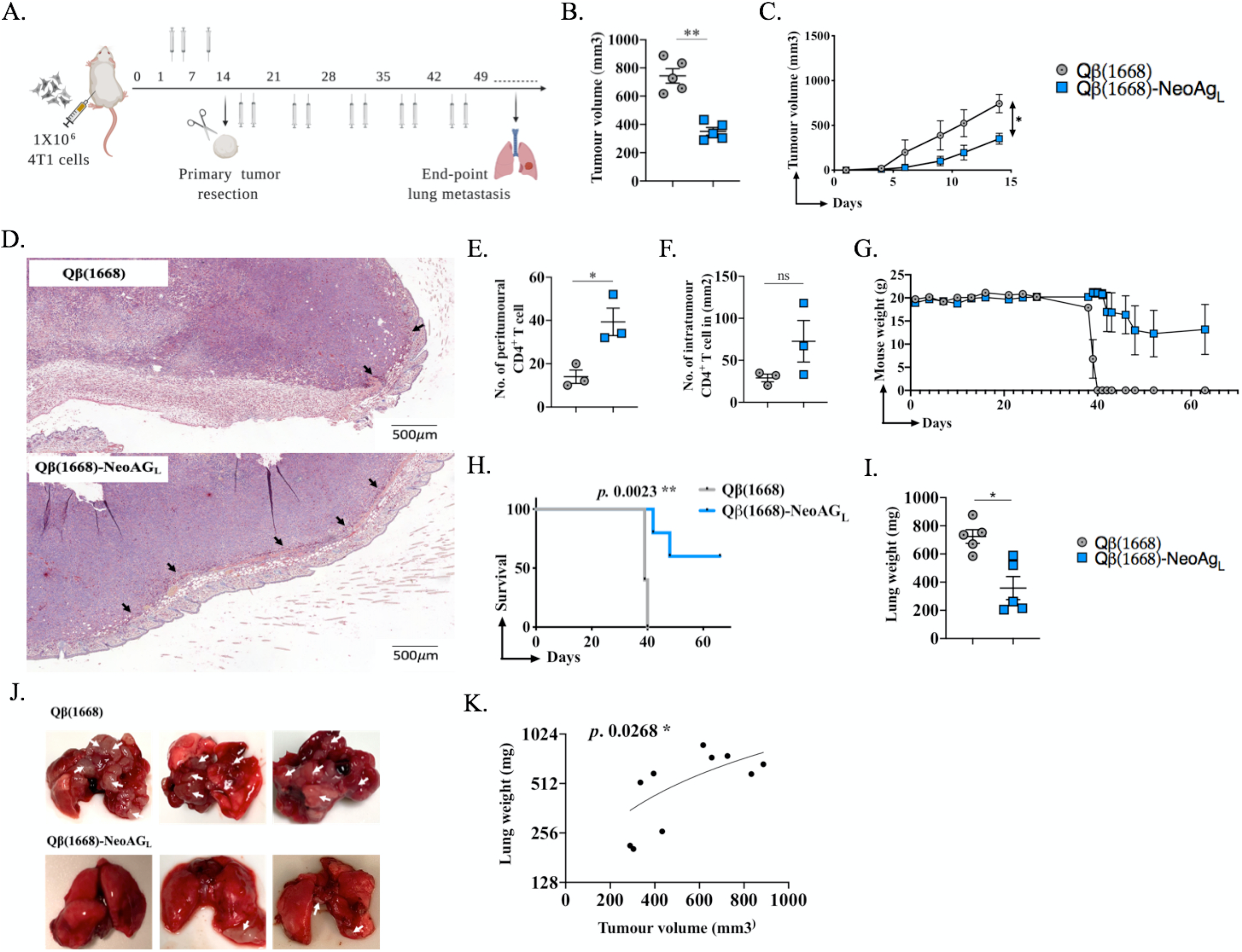
Long peptide vaccination decreases post-surgical tumour recurrence and lung metastases, and prolongs mouse survival. *A*, Mice were inoculated with 1×10^6^ 4T1 cells on day 0 and vaccinated 3 times over 13 days. Primary tumours were surgically resected on day 14 under isoflurane anaesthesia, followed by 2 vaccinations weekly until the end of the experiment. Mice were followed daily to monitor tumour recurrence, general health and mouse weight. *B*, Tumour volume in (mm3) (means ± SEM) measured on day 14 post 4T1 inoculation of control mice treated with Qβ(1668) or mice vaccinated with Qβ(1668)-NeoAG_L_. Each dot represents an individual tumour. *C*, Tumour growth curves of the treated groups, AUC was measured and statistical analysis was done by Student’s *t*-test (2-tailed). *D*, Representative immunohistochemistry staining of CD4^+^ T cells in tumours of mice treated with Qβ(1668) as control or Qβ(1668)-NeoAG_L_. Arrows point to peritumoural CD4^+^ T cells (n=3). *E*, Numbers of peritumoural and intratumoural/mm2 CD4^+^ T cells, respectively. *G*, Mice weight (g) during the experiment. *H*, Survival of mice in both groups; mice were euthanized when loosing 10-15% of their original weight, statistical analysis by log-rank test. *I*, Lung weight (mg), mice were euthanized when loosing 10-15% of their original weight and lungs were collected. *J*, Representative photos of lungs from euthanized mice (control and Qβ(1668)-NeoAG_L_); arrows point to lung metastases. For the group treated with Qβ(1668)-NeoAG_L_, two lung samples are from the mice which were euthanized due to weight loss and one lung from a mouse that remained healthy and without metastases until the end of the experiment. *K*, correlation between primary tumour volume in (mm3) and end-point lung weight (mg). Statistical analysis by Student’s *t*-test. (n=5). P values \*\*\*\**P <* 0.0001; \*\*\**P <* 0.001; \*\**P <* 0.01; \**P <* 0.05.

## Discussion

In this study, we developed a universal approach for a personalized vaccine. We found efficient tumour control through activation and functional competence of both CD8^+^ and CD4^+^ T cells, associated with reduction of suppressive myeloid cells both in circulation and within tumours. The vaccine was based on the successful identification of neoantigenic peptides using mass spectrometry-based immunopeptidomics combined with whole exome sequencing and wet-bench validation. The advantage of this experimental approach relies on the fact that it only identifies peptides that are indeed naturally presented by MHC molecules on the surface of tumour cells, thus avoiding the frequent epitope prediction failures by other techniques that do not rely on the physical identification of peptides. Indeed, our approach has been shown to enrich for neoantigens that are capable of controlling tumours in mice *in vivo* (41). To avoid possible outgrowth of tumour escape variants, we produced a multi-target vaccine with four neoantigenic peptides.

We synthesized short peptides corresponding to CD8^+^ T-cell epitopes or long peptides including both CD8^+^ and CD4^+^ T-cell epitopes. Using the bio-orthogonal Cu-free click chemistry, peptides were coupled to VLPs consisting of recombinant bacteriophage Qβ protein and loaded with the B-type CpG 1668. The applied chemical coupling has several advantages as compared to other methods such as using a SMPH linker (42). SMPH may react with internal cysteine epitopes, causing their inactivation. Additionally, non-reacted SMPH on VLPs can be toxic and may complicate the GMP process. Cu-free click chemistry is a non-toxic method, as the azide group added to the C-terminal end of the synthesized peptides is metabolically stable and lack any reactivity with other biological functionalities in human cells. Cu-free chemistry has shown high selectivity in several studies, for example for imaging, labelling and tracking cells *in vivo* (43, 44). A key advantage of this method is that it can be applied at the bedside, enabling easy and rapid production of personalized vaccines for each patient directly where and when needed.

The short peptide vaccine significantly hindered tumour progression (*p*. 0.0133) when compared to the antigen negative control vaccine. However, this vaccine did not enhance CD8^+^ nor CD4^+^ T-cell infiltration into the tumours. Nevertheless, it significantly decreased (*p*. 0.0132) circulating MDSCs. The insufficient anti-tumour efficacy of the short vaccine may be due to deficiency in activating CD4^+^ helper T cells that are often required to support the CD8^+^ T cells as shown in several studies (8, 9, 45, 46). Indeed, vaccination with elongated peptides (by addition of natural flanking sequences) significantly increased the anti-tumour efficacy (*p*. 0.0091). Additionally, long peptide vaccination elicited stronger T-cell responses characterized by increased percentages of polyfunctional specific CD8^+^ and CD4^+^ T cells. Furthermore, long peptide vaccination enhanced the infiltration of CD4^+^ T cells into the tumours. It is known that CD4^+^ T cells can license DCs via CD40-CD40L interaction which enhances antigen presentation to and activation of T cells (47). It may also be possible that CD4^+^ T cells themselves kill tumour cells (45, 48). Probably more important is that tumour growth is inhibited by pro-inflammatory milieu generated by the effector cytokines IFN-γ and TNF-α which are not only released by CD8^+^ T cells but also by CD4^+^ T cells.

There is general agreement that T-cell vaccines must become more potent, particularly for the induction of CD8^+^ T-cell responses, in order to increase the likelihood of clinical benefit. However, while CD4^+^ T-cell responses are readily induced by the currently used cancer vaccines (e.g. RNA, or free peptides mixed with adjuvants), CD8^+^ T-cell responses induced by the majority of vaccines remain relatively scarce (4, 5), particularly when vaccine antigens must undergo processing and cross-presentation by DCs which is an important but relatively inefficient process. In turn, the requirement for cross-presentation has the advantage that only DCs present the vaccine’s MHC-I antigens to CD8^+^ T cells, favouring robust T-cell priming and boosting, and avoiding tolerance induction (49).

Immunosuppressive mechanisms are often responsible for the inefficiency of immunotherapy. Some of those mechanisms may be successfully hindered by therapies that appropriately activate T-cell responses. Indeed, our long peptide vaccine led to significant reduction within tumours of granulocytic and myeloid-derived suppressor cells (MDSCs) known to exert negative effects on T cells, whereas appropriately activated T cells produce IFN-γ that counteracts MDSCs (50). These two cell populations may also be used as biomarkers for predicting treatment efficacy.

Surgical resection is usually the first step in the clinical management of cancer. Subsequently, residual microtumours as well as circulating cancer cells may cause potentially lethal recurrence (51). Immunotherapy is particularly necessary and potentially useful in patients who are at risk for subsequent disease recurrence and progression. Checkpoint blockade is often used in this setting. Alternatively, or in addition, personalized cancer vaccines may be considered, as they have the potential to activate the immune system more specifically and may thus have a favourable efficacy/toxicity profile, provided that they achieve sufficient anti-tumour effects for providing clinical benefit. This can be well addressed in the 4T1 tumour model, because these tumours can spontaneously metastasize from primary lesions to lymph nodes, lung, brain and bone, resembling triple-negative stage IV human breast cancer (52). After surgical resection of the primary tumours, we found that our multi-neoantigen vaccine led to reduction of tumour recurrence and metastases formation. Therefore, the use of our vaccine in this setting may be promising.

Vaccination alone may be insufficient to achieve satisfactory tumour control (53) calling for combination therapies. For example, we and others have shown previously that combining a multi-targeting vaccine with anti-CD25 mAb treatment to inhibit regulatory T cells significantly improved the anti-tumour efficacy in challenging mouse tumours (22, 54). We envision that combining our new vaccine with anti-CD25 mAb, anti-OX40 mAb and/or checkpoint inhibitors may further improve therapy efficacy. The identification of patient-individual neoantigens remains challenging but is technically feasible (28, 55). Together, our approach represents a further step making the vision of personalized cancer vaccines more realistic. Clinical development is encouraged because the vaccine induces efficient anti-tumour responses and is practically feasible as it can be formulated at the bedside and thus widely applied.

## Acknowledgment

This work was supported by Qatar National Research Fund (PDRA4-0118-18002) and Swiss Cancer Research (KFS-4291-08-2017-R). The graphical abstract, Figure 1A, Figure 3A and Figure 5A were generated with BioRender. Open Access funding provided by the Qatar National Library

## Authors Contributions

Design of experiments, acquisition of data, interpretation and analysis of data: MOM, DES, SdB and MFB. Whole exome sequencing and mass spectrometry-based identification of neoantigens: MBS, JM, HP, BJS, GC. Writing, revision and editing of manuscript: MOM, DES, MBS and MFB. Technical, material and tool support: VI, MV, SD, PS. Study supervision: MOM and MFB. All authors read and approved the final manuscript.

---

**Supplementary Figure 1. Sequences of neoantigens and corresponding non-mutated antigens and verification by RT-RCR.** A, (Top) NeoAG1 (E>K) (KYLDSPKRL) from 4T1 mammary carcinoma. A, (Bottom) Corresponding WT sequence (KYLDSPERL) from EpH4-Ev control mammary cell-line. B, (Top) NeoAG2 (E>K) (SPKRLVGL) from 4T1 mammary carcinoma. B, (Bottom) Corresponding WT sequence (SPERLVGL) from EpH4-Ev control mammary cell-line. C, (Top) NeoAG3 (H>Y) (KYNAVPTVF) from 4T1 mammary carcinoma. C, (Bottom) Corresponding WT sequence (KHNAVPTVF) from EpH4-Ev control mammary cell-line. D, (Top) NeoAG4 (H>N) (VYREQMNGV) from 4T1 mammary carcinoma. D, (Bottom) Corresponding WT sequence (VYREQMHGV) from EpH4-Ev control mammary cell-line.

**Supplementary Figure 2. Representative immunohistochemistry staining of CD4^+^ T cells for tumours treated with Qβ(1668) as a control or Qβ(1668)-NeoAG_L_.** Arrows point to intratumoural CD4^+^ T cells. 3 samples per group were analysed.

**Supplementary Table 1. MHC-I and MHC-II binding peptides that include the mutations detected by peptidomics.** MHC ligand predictions were performed with NetMHCpan tool (MHC allele and % rank score). MaxQuant identification score is indicated for the neoantigens detected my MSMS. Bold, mutated residues; red, lysine residues required for the click chemistry.

**Supplementary Data 1**. 4T1 derived MHC-I peptides identified by MSMS (Excel Sheet)

